# GABA AND GLUTAMATE CHANGES IN PEDIATRIC MIGRAINE

**DOI:** 10.1101/2020.04.14.041616

**Authors:** Tiffany Bell, Mehak Stokoe, Akashroop Khaira, Megan Webb, Melanie Noel, Farnaz Amoozegar, Ashley D Harris

**Affiliations:** Department of Radiology, University of Calgary, Calgary, AB, Canada; Hotchkiss Brain Institute, University of Calgary, Calgary, AB, Canada; Alberta Children’s Hospital Research Institute, University of Calgary, Calgary, AB, Canada; Department of Psychology, University of Calgary, Calgary, AB, Canada; Department of Clinical Neurosciences, University of Calgary, Calgary, AB, Canada

## Abstract

Despite migraine being one of the top five most prevalent childhood diseases, a lack of knowledge about pediatric migraine limits effective treatment strategies; standard adult pharmaceutical therapies are less effective in children and can carry undesirable side-effects. Non-pharmacological therapies have shown some success in adults; however, to appropriately apply these in children we need to understand pediatric migraine’s underlying biology. One theory is that migraine results from an imbalance in cortical excitability. Magnetic resonance spectroscopy (MRS) studies show changes in GABA and glutamate levels (the primary inhibitory and excitatory neurotransmitters in the brain, respectively) in multiple brain regions. Although there is indirect evidence of abnormal excitability in pediatric migraine, GABA and glutamate levels have yet to be assessed.

The purpose of this study was to measure levels of GABA and glutamate in the thalamus, sensorimotor cortex and visual cortex of children with migraine using MRS. We found that children with migraine and aura had significantly lower glutamate levels in the visual cortex as compared to control children, opposite to results seen in adults. Additionally, we found significant correlations between metabolite levels and migraine characteristics; higher GABA levels were associated with a higher migraine burden. We also found that higher glutamate in the thalamus and higher GABA/Glx ratios in the sensorimotor cortex were associated with duration since diagnosis, i.e., having migraines longer. Lower GABA levels in the sensorimotor cortex were associated with being closer to their next migraine attack. Together this indicates that GABA and glutamate disturbances occur early in migraine pathophysiology and emphasizes that evidence from adults with migraine cannot be immediately translated to paediatric sufferers. This highlights the need for further mechanistic studies of migraine in children, to aid in the development of more effective treatments.

## INTRODUCTION

Migraine often begins in childhood, with roughly 20% of sufferers experiencing their first attack before the age of 5.^1^ Early intervention can decrease migraine frequency, with those receiving earlier interventions more likely to achieve remission.^2^ However, treatment strategies for children are limited, in part due to limited knowledge about pediatric migraine biology.^3^ Migraine is often managed similarly in children as adults, despite evidence that children with migraine present with different symptoms.^4^ Standard adult medications for preventing migraine have been shown to be no more effective than placebo in children, and often come with undesirable side effects.^1,4,5^ Non-pharmacological therapies such as brain stimulation^6^ and neurofeedback^7^ have shown some success in adults; however, to appropriately adapt these for children we need to understand the underlying biology of pediatric migraine.

There is increasing evidence that adult migraine results from an imbalance of excitation/inhibition in the brain, which changes cyclically until a migraine occurs (known as the migraine cycle).^8,9^ During the interictal period, excitability of the cortex increases proportionally with time to the next attack. During the migraine or shortly thereafter, the brain returns back to baseline activity and begins the cycle again.^8,9^ Neurophysiological studies suggest this cortical hyperexcitability is a result of abnormal thalamic control.^10^ Studies have revealed that, in migraine, there is altered communication in the thalamo-cortical networks^11^ which underlie important processes in multisensory integration; this altered communication is associated with clinical migraine symptoms.^10,12^ The excitability level of the sensory cortices is set by activity in these thalamo-cortical loops. Between attacks, adults with migraine have shown reduced activity of early high frequency oscillation bursts during somatosensory evoked potentials, indicating reduced function in thalamocortical connectivity.^13^ This results in increased excitability in the sensory and visual cortices. For example, the sensorimotor cortex of migraineurs shows enhanced responses to sensory stimuli, and the degree of enhancement correlates with headache frequency.^14^ Similarly, adults with migraine have lower thresholds for transcranial magnetic stimulation (TMS) induced visual phosphenes (i.e., a ring of light in the participant’s vision) compared to controls, indicating visual cortex hyperexcitability.^15^

The primary neurochemicals associated with inhibition and excitation in the human brain are GABA and glutamate, respectively. These can be measured *in vivo* non-invasively using magnetic resonance spectroscopy (MRS). A recent review showed that, in adults with migraine, there is an increase in GABA and glutamate levels measured using MRS in multiple brain areas.^16^ For example, adults with migraine show increased glutamate^17–20^ and increased GABA levels^21,22^ in the visual cortex, and increased glutamate in the thalamus.^20^

In children with migraine, there is also evidence of excitation/inhibition imbalances. Studies have shown children with migraine have shortened somatosensory evoked potential recovery cycles, suggesting disinhibition at different levels of the central nervous system.^23^ Consistent to what is observed in adults, children with migraine also have lower phosphene thresholds which increase before a migraine attack,^24^ demonstrating changes in excitability of the visual cortex.

Despite evidence of abnormal cortical excitability in children with migraine, GABA and glutamate levels remain uninvestigated. Additionally, the GABA/Glutamate ratio may be used to index the inhibitory/excitatory balance and may show a stronger effect than changes in either GABA or glutamate alone. The aim of the present study was to use MRS measures to investigate GABA and glutamate in the thalamus, sensorimotor and visual cortices of children with migraine. We expected children with migraine to show higher levels of GABA and glutamate in all three brain regions, as has been observed in the visual cortex of adults with migraine.

## MATERIALS AND METHODS

### Ethics Statement

The study protocol was approved by the Conjoint Health Research Ethics Board (CHREB), University of Calgary. All of the study participants provided informed assent and their parents provided informed consent at time of enrollment.

### Participants

35 children with migraine (Migraine group) aged 7-13 years with a diagnosis of migraine from their family physician were recruited from the Vi Riddell Pain Clinic at the Alberta Children’s Hospital and the local community. Participants were included if they had received a physician diagnosis of migraine, which was confirmed using the ICHD-III beta diagnostic criteria,^25^ with no other accompanying neurological, psychiatric or systematic disorders (e.g. ADHD, Autism), they met the standard MRI safety criteria (e.g. no metal implants or devices) and were not taking preventative medications such as triptans.

31 age and sex-matched children without migraine (Control group) were recruited using the Healthy Infants and Children Clinical Research Program (HICCUP). The same exclusion criteria were applied (no neurological, psychiatric or systemic disorders and standard MRI safety criteria), with the exception of migraine history: volunteers were included if they had no history of migraine.

### Migraine Diary

The parents of children with migraine were asked to keep a migraine diary for 30 days preceding their appointment and 7 days following, which was sent to their computer or mobile device. In this diary, they were asked to record if their child had had a migraine that day and, if so, the length and pain level of the attack (using the Wong-Baker FACES pain rating scale26 of 1-10), along with how they treated the migraine. If the child did not have a migraine in the 7 days following the appointment, they were asked to provide the date of the following migraine. Using the migraine diaries, the *“position in the migraine cycle”* at the time of scanning was calculated as the number of days since the last migraine divided by the number of days between the last migraine and the next. A higher number represents being further along in the cycle and, subsequently, closer to the next migraine, while accounting for inter-individual differences in overall length of the migraine cycle. Children with migraine were excluded if they did not experience a migraine in the 30 days preceding their appointment, or if they experienced a migraine on the day of the appointment.

### Questionnaires

All participants completed the following questionnaires: Headache Impact Test (HIT-6),^27^ Pediatric Migraine Disability Assessment (PedMIDAS),^28^ Revised Children’s Anxiety and Depression Scale (RCADS) short version,^29^ Puberty Status Scale^30^ and the Edingburgh Handedness Scale.^31^ The HIT-6 is a 6-item self report survey that assesses the negative impact of headaches on normal daily activity. Responses range from 36-78, with higher scores representing more negative impact. PedMIDAS is a pediatric version of the self report Migraine Disability Assessment (MIDAS) questionnaire commonly used in adults, and has been validated in children and adolescents ranging from 6-18.^28^ Responses range from 0-90 with higher scores representing more negative impact. The RCADS is a self report measure developed to assess anxiety and depression symptoms among children and adolescents and has been validated in children and adolescents ranging from 6-18.^32^ Responses range from 0-45 for anxiety and 0-30 for depression, with higher scores representing more symptoms. The Pubertal Status Scale is a self-report measure based on the Tanner pubertal staging, scores are categorised into the following stages: prepubertal, early pubertal, mid pubertal, late pubertal and post pubertal.^30^

### MRI Acquisition

Scanning was performed on a 3T GE 750w MR scanner using a 32-channel head coil. A T1-weighted anatomical image was collected for voxel placement (BRAVO; TE/TR = 2.7/7.4ms, slice = 1 mm^3^ isotropic voxels). 3×3×3 cm^3^ voxels were placed in the thalamus (midline centred), right sensorimotor cortex (centred on the hand-knob of the motor cortex, and rotated such that the coronal and sagittal planes aligned with the cortical surface^33^) and the occipital cortex (as close to aligning with the parietoocciptal sulcus as possible, without including cerebellum, midline centred; Figure 1).

**Figure 1:**
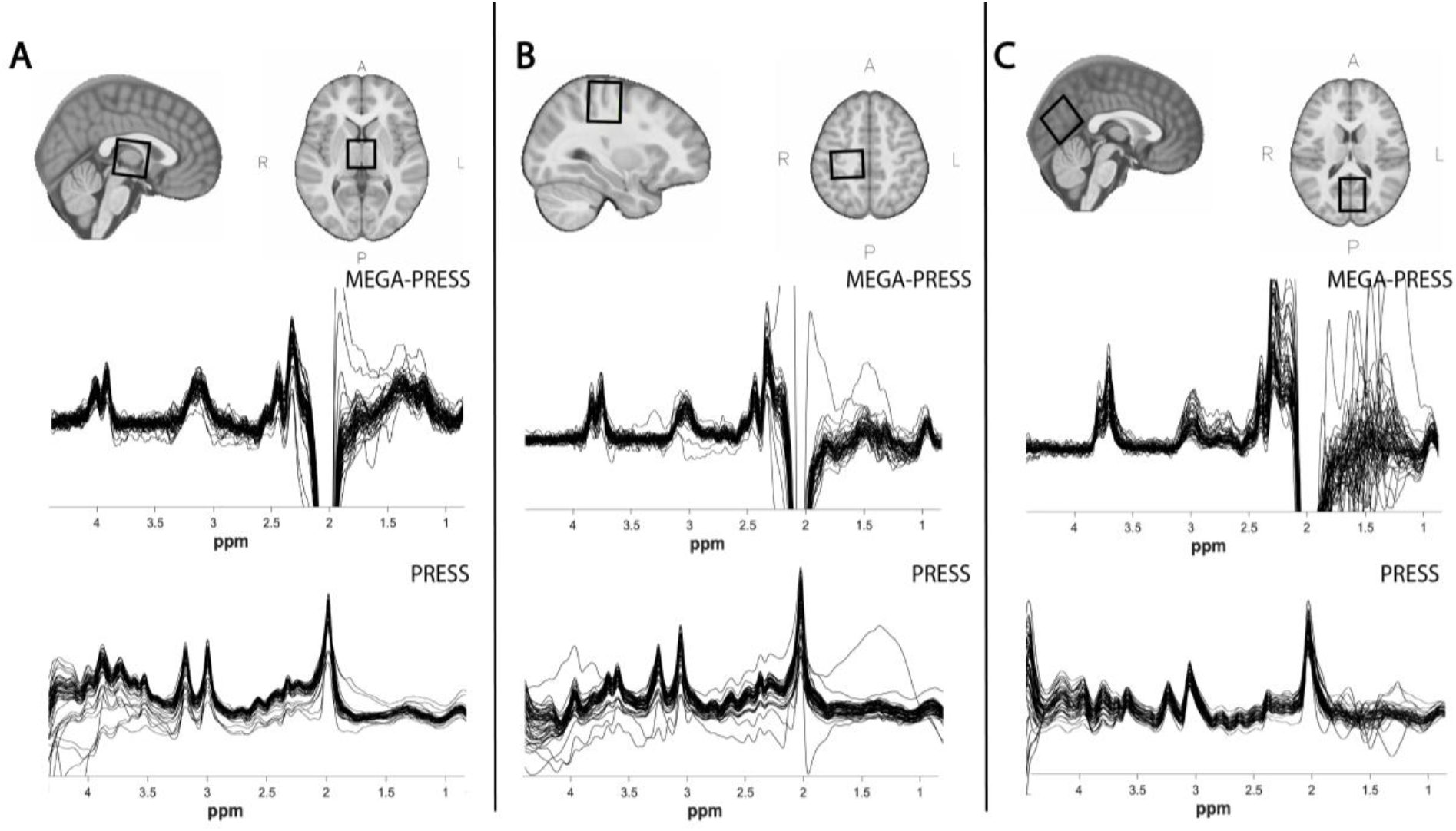
Example voxel placement (top row) and overlay of all (migraine and control) MEGA-PRESS (middle row) and PRESS spectra (bottom row) for A) Thalamus, B) Sensorimotor Cortex, C) Visual Cortex.

GABA-edited spectroscopy data were collected using macromolecule suppressed MEGA-PRESS (TR/TE = 1800/80 ms, 20 ms editing pulses at 1.9 and 1.5 ppm, 256 averages) from each brain area. Separate PRESS data (TR/TE = 1800/35 ms, 64 averages) were also acquired from each region to quantify glutamate.

### MRI Analysis

MEGA-PRESS data were analysed using Gannet3.1,^8^ which included the following preprocessing steps: coil combination, frequency and phase correction, apodization and down-weighting of motion-corrupted averages. Tissue correction was performed using voxel tissue fractions obtained by generating a subject-specific voxel mask registered to each individual tissue segmented T1 anatomical image.^9^

PRESS data were preprocessed with the FID-A^36^ toolbox using the following preprocessing steps: coil combination, removal of motion-corrupted averages, frequency drift correction and zero order phase correction.^37^ LCModel Version 6.3-1J^38^ was used to apply eddy current correction and quantification relative to water. Basis sets for quantification (including alanine, aspartate, glycerophosphocholine, phosphocholine, creatine, phosphocreatine, GABA, Glu, Gln, lactate, inositol, n-acetyl aspartate, n-acetylaspartylglutamate, scyllo-inositol, glutathione, glucose and taurine) were simulated using the FID-A toolbox based on exact sequence timings and RF pulse shapes. Metabolite values were corrected for tissue composition^39^ using the tissue fractions generated from Gannet3.1. Glutamate was quantified both on its own (Glu) and as a combination (Glx) of glutamate and it’s precursor, glutamine. Glutamate and glutamine overlap on the spectra due to their similar chemical compositions, subsequently it is difficult to separate the individual signals. Data quality was assessed by visual inspection and metabolite linewidth, spectra with a linewidth over 0.1 ppm were excluded.

### Statistical Analysis

ANCOVAs were used to compare metabolite levels between groups, controlling for age. Correlation analyses, also controlling for age, were used to test the relationship between metabolite levels and migraine characteristics.

## RESULTS

### Participants

Participant demographic data and questionnaire results are shown in Table 1, the final sample sizes were 29 Migraine and 27 Control. From the 35 children with migraine recruited, 2 children were excluded as they did not experience a migraine during the past 30 days and 3 children were excluded as they experienced a migraine on the day of the scan. Imaging data was not acquired from 1 child with migraine. From the 31 age and sex-matched controls recruited, 1 child was excluded as they developed migraines shortly after participating in the study, 1 child was excluded due to an ADHD diagnosis, and imaging data was not acquired for two control children. Some MRS data was not collected due to children requesting to get out of the scanner and individual voxel data of poor quality was removed on a case-by-case basis. Table 2 shows the final number of spectra for each voxel included for each analysis. There were no significant differences in data quality (measured using linewidth) between the two groups.

**Table 1:** Table of demographics and questionnaire results (mean ± SD).

|  |  | Migraine | Control | Comparison |
| --- | --- | --- | --- | --- |
| **N** | | 29 (12 female) | 27 (14 female) | $X^2(1,57)=0.414$, $p=0.600$ |
| **Age (years)** | | 10.19 (1.3) | 9.9 (1.5) | $t(55)=-1.009$, $p=0.317$ |
| **HIT-6** |  | **59.07 (7.2)** | **43.5 (4.8)** | **$F(1,53)=80.54$, $p<0.001$** |
| **PedMIDAS** |  | **21.5 (19.8)** | **2.0 (3.6)** | **$F(1,54)=24.87$, $p<0.001$** |
| **RCADS Depression** | | 7.6 (4.5) | 6.1 (4.2) | $F(1,54)=1.754$, $p=0.191$ |
| **RCADS Anxiety** | | 10.6 (7.5) | 7.5 (5.1) | $F(1,54)=3.884$, $p=0.053$ |
| **Pubertal Status (N)** | **Pre** | 18 | 10 | $X^2(3,57)=4.488$, $p=0.213$ |
|  | **Early** | 7 | 10 |  |
|  | **Mid** | 4 | 7 |  |
| **Number of Migraines in 30 days** |  | 6.3 (6.9) | N/A | N/A |
| **Range** |  | 1-30 | N/A | N/A |
| **Disease Duration (years)** |  | 3.9 (2.5) | N/A | N/A |

**Table 2:**
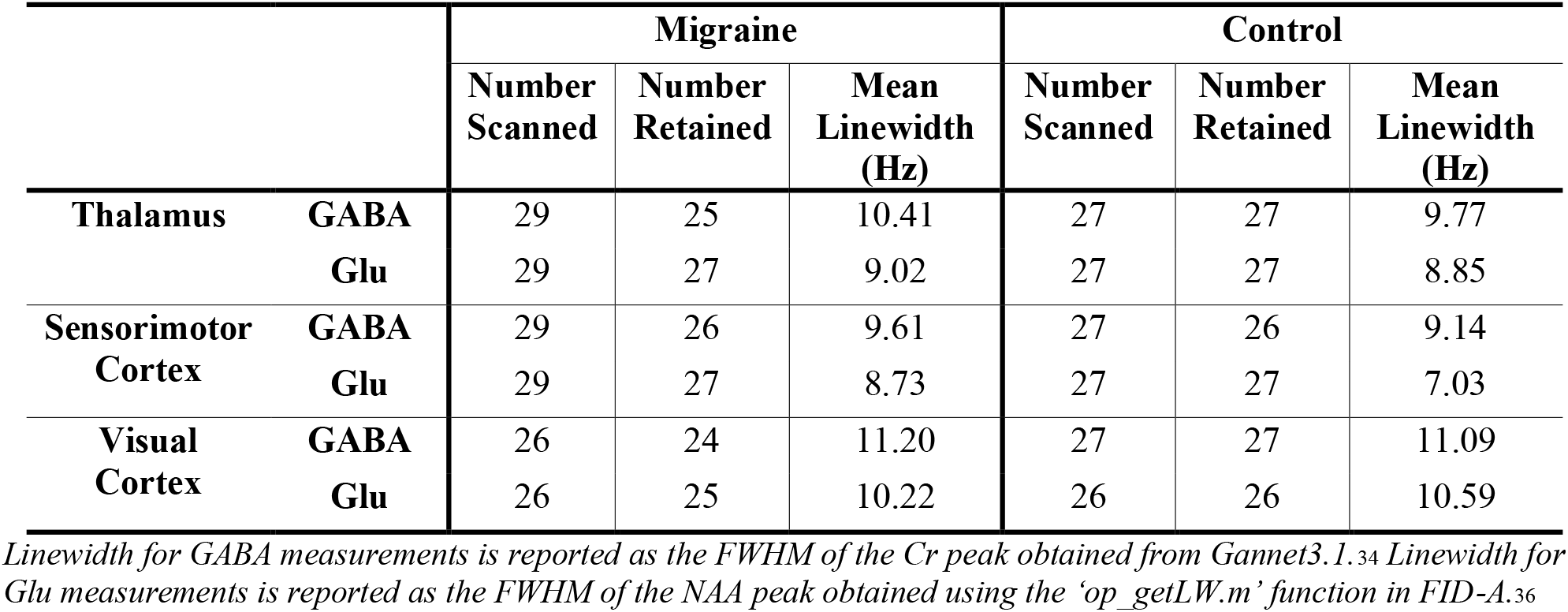
Table showing number of participants included for each brain region analysis, and group mean linewidth

|  |  | Migraine |  |  | Control |  |  |
| --- | --- | --- | --- | --- | --- | --- | --- |
|  |  | Number Scanned | Number Retained | Mean Linewidth (Hz) | Number Scanned | Number Retained | Mean Linewidth (Hz) |
| Thalamus | GABA | 29 | 25 | 10.41 | 27 | 27 | 9.77 |
|  | Glu | 29 | 27 | 9.02 | 27 | 27 | 8.85 |
| Sensorimotor Cortex | GABA | 29 | 26 | 9.61 | 27 | 26 | 9.14 |
|  | Glu | 29 | 27 | 8.73 | 27 | 27 | 7.03 |
| Visual Cortex | GABA | 26 | 24 | 11.20 | 27 | 27 | 11.09 |
|  | Glu | 26 | 25 | 10.22 | 26 | 26 | 10.59 |

Linewidth for GABA measurements is reported as the FWHM of the Cr peak obtained from Gannet3.1.<sup>34</sup> Linewidth for Glu measurements is reported as the FWHM of the NAA peak obtained using the 'op\_getLW.m' function in FID-A.<sup>36</sup>

### Group Comparisons

#### Thalamus

There were no significant differences in Glx (F(1,51)=0.241, p=0.626), Glu (F(1,51)=0.001, p=0.974) or GABA (F(1,49)=0.431, p=0.515) levels in the thalamus between the two groups. There was a trend towards higher GABA/Glx ratios in Migraine (F(1,48)=2.950, p=0.092), mainly driven by higher GABA levels (

#### Sensorimotor Cortex

There were no significant differences between groups in any of the metabolites in the sensorimotor cortex (Glx: F(1,50)=0.870, p=0.356; Glu: F(1,50)=1.047, p=0.311; GABA: F(1,49)=0.927, p=0.340; GABA/Glx: F(1,47)=1.258, p=0.268; Figure 3).

**Figure 2:**
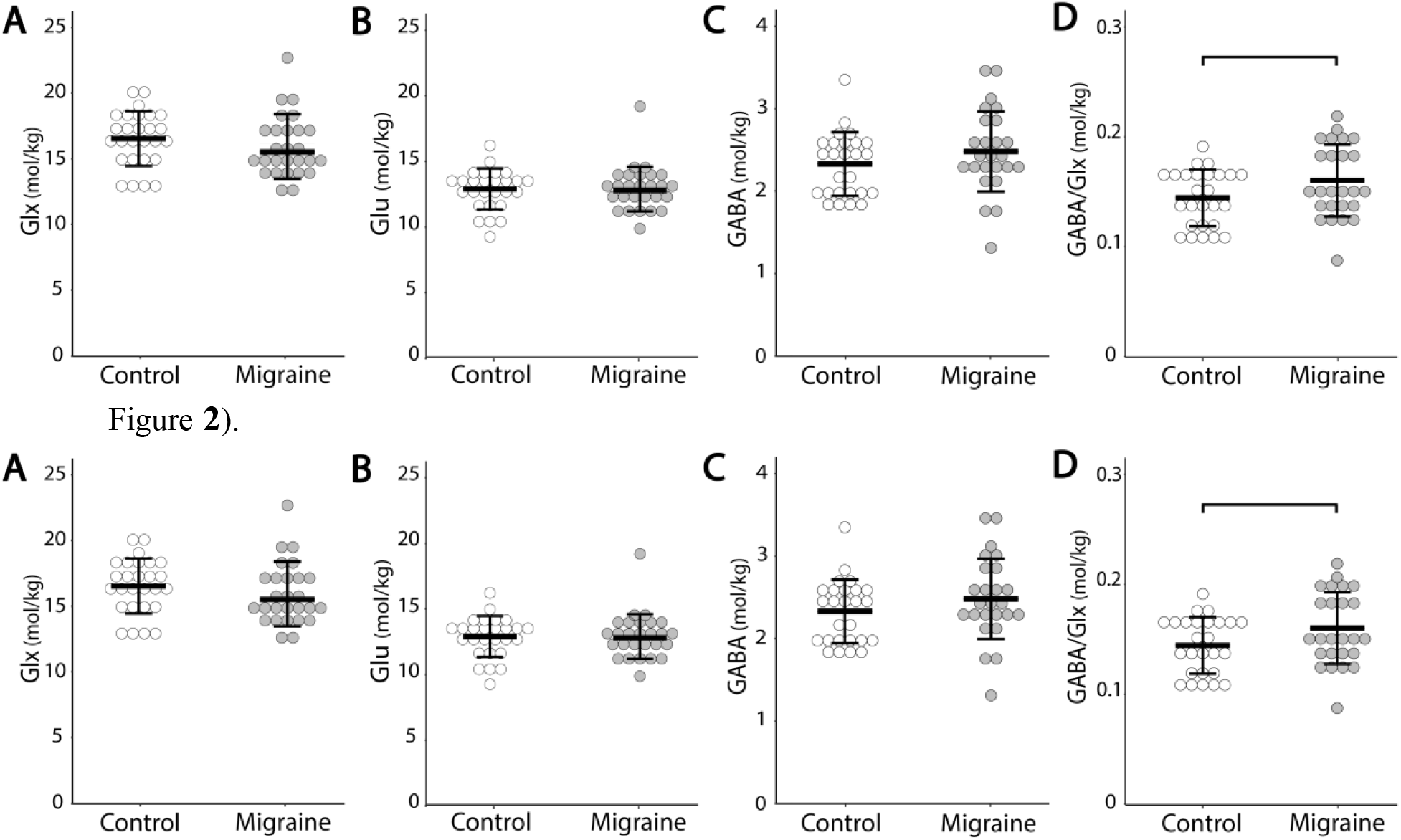
Group comparisons of **A**: Glx levels (F(1,51)=0.241, p=0.626); **B**: Glu levels (F(1,51)=0.001, p=0.974); **C**: GABA levels (FF(1,49)=0.431, p=0.515); **D**: GABA/Glx ratios (FF(1,48)=2.950, p=0.092) in the thalamus. Each circle represents an individual subject.

**Figure 3:**
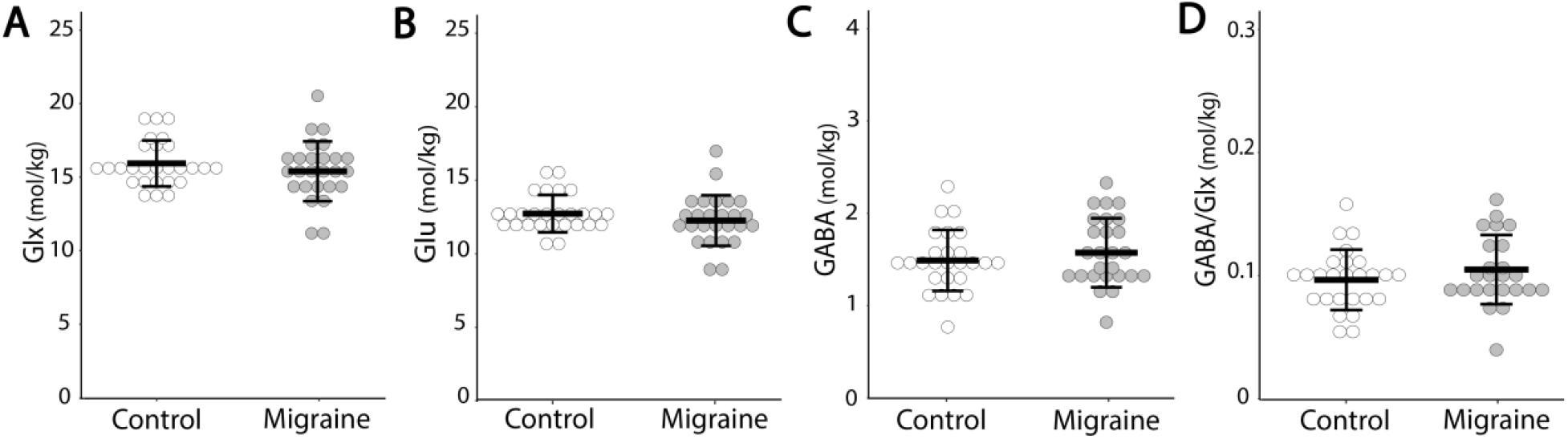
Group comparisons of **A**: Glx levels (F(1,50)=0.870, p=0.356); **B**: Glu levels (F(1,50)=1.047, p=0.311); **C:** GABA levels (F(1,49)=0.927, p=0.340); **D**: GABA/Glx ratios (F(1,47)=1.258, p=0.268) in the sensorimotor cortex. Each circle represents an individual subject.

#### Visual Cortex

There were no significant differences in Glx (F(1,48)=1.989, p=0.165), Glu (F(1,48)=2.812, p=0.100), GABA (F(1,48)=0.582, p=0.449), or GABA/Glx ratios (F(1,47)=1.276, p=0.264; Figure 4) in the visual cortex between the two groups.

**Figure 4:**
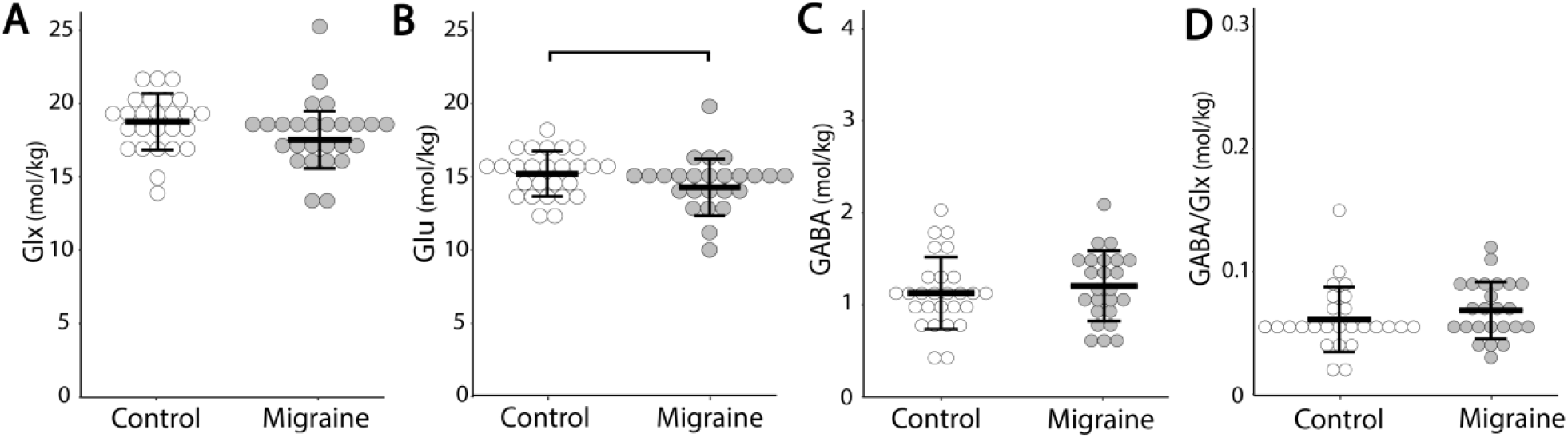
Group comparisons in **A**: Glx levels (F(1,48)=1.989, p=0.165); **B**: Glu levels (F(1,48)=2.812, p=0.100); **C**:GABA levels (F(1,48)=0.582, p=0.449); **D**: GABA/Glx ratios (F(1,47)=1.276, p=0.264) in the visual cortex. Each circle represents an individual subject.

##### Aura

Of the 25 children with migraine included in the visual cortex analyses, 9 reported aura either before or during their migraine, 15 reported no aura and 1 participant did not specify. There was a significant effect of group on Glu levels (F(2,46)=3.317, p=0.045) when dividing the migraine group into aura or non-aura. Levels of Glu were significantly lower in Migraine with aura (p=0.022) compared to Controls. There were no significant differences in GABA (F(2,46)=0.458, p=0.635), Glx (F(2,46)=2.798, p=0.071) or GABA/Glx ratios (F(2,45)=1.024, p=0.367) in the visual cortex between the three groups.

### Migraine Characteristics

In Migraine, higher glutamate levels in the thalamus (r(17)=0.514, p=0.025, Figure 6A) and higher GABA/Glx ratios in the sensorimotor cortex (r(15)=−0.562, p=0.019, Figure 6B) were associated with a greater number of years with migraine.

**Figure 5:**
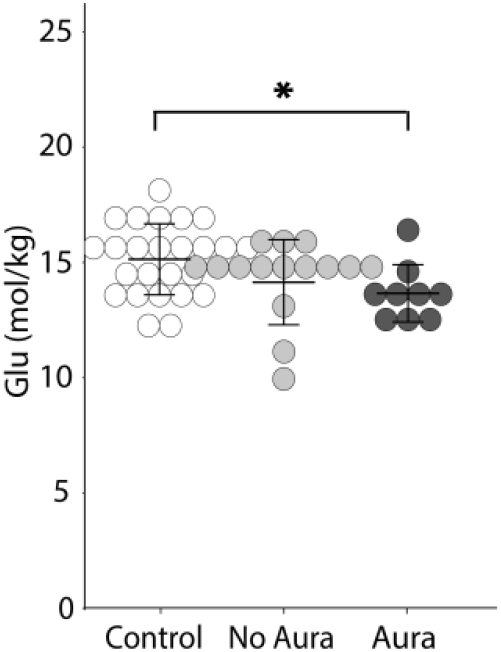
Glu levels in Migraine with and without aura, and Controls. Migraine with aura had significantly lower glutamate levels in the visual cortex compared to Control (p=0.019). There was no significant difference between Migraine without aura and control (p=0.135) or between the Migraine with and without aura groups (p=0.324).

**Figure 6:**
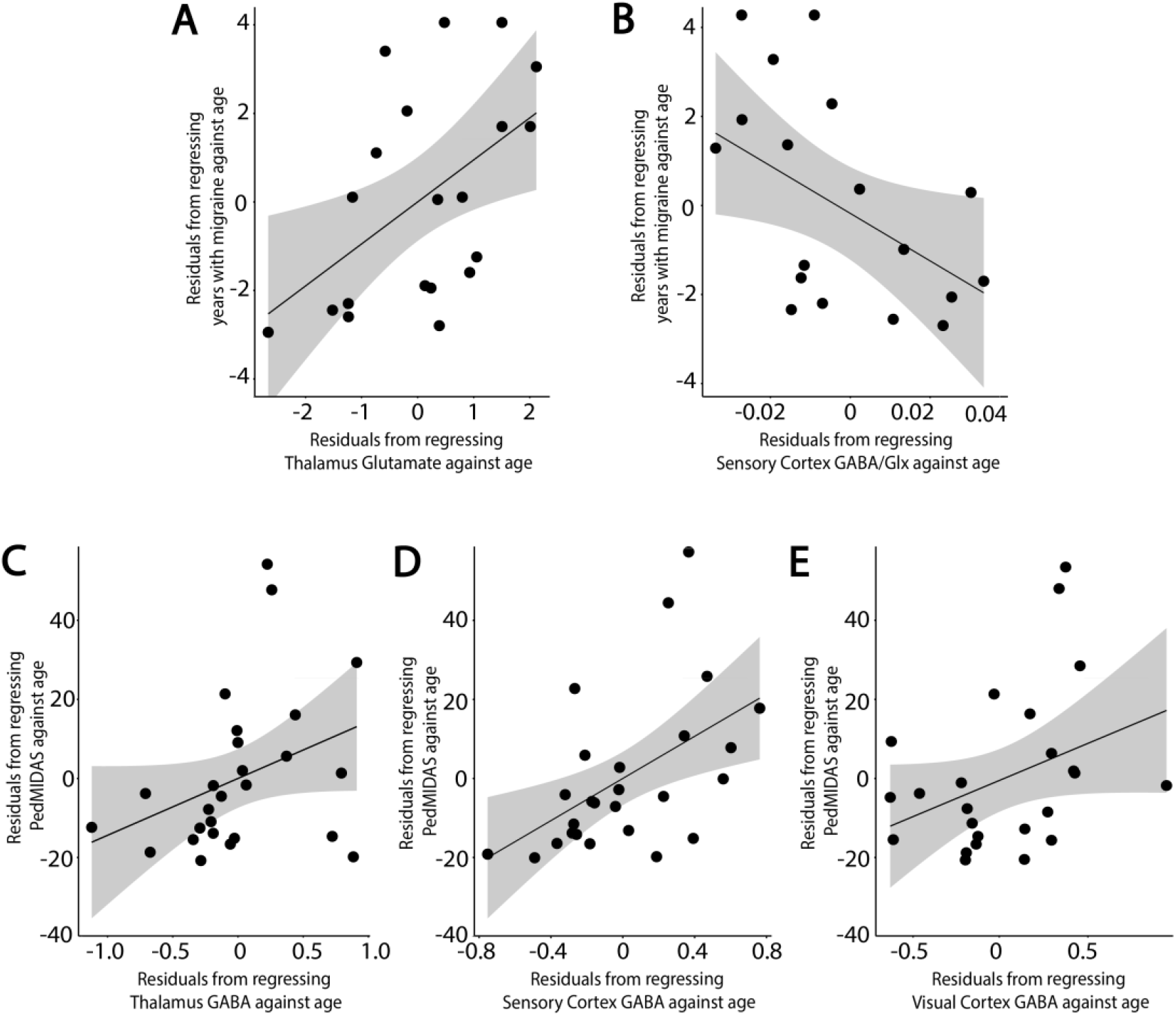
Associations between migraine characteristics and neurochemical levels. **A**: Positive correlation between Glu in the thalamus and the number of years with migraine, controlling for age (r(18)=0.532, p=0.016); **B**: Negative correlation between the GABA/Glx ratio in the thalamus and the number of years with migraine, controlling for age (r(17)=−0.456, p=0.05); **C**: Positive association between GABA in the thalamus and the PedMIDAS score, controlling for age (r(23)=0.370, p=0.068); **D**: Positive association between GABA in the sensorimotor and the PedMIDAS, controlling for age (r(24)=0.514, p=0.007); **E**: Positive association between GABA in the visual cortex and the PedMIDAS, controlling for age (r(22)=0.361, p=0.084). Grey shading indicates the 95% confidence intervals on the partial correlations.

In Migraine, higher levels of GABA in the sensorimotor cortex were associated with higher PedMIDAS scores (r(23)=0.516, p=0.008, Figure 6D), indicating children with higher GABA levels were more impacted by their migraines. This association was also seen in the thalamus and visual cortex but did not reach statistical significance. (Thalamus: r(22)=0.365, p=0.080, Visual Cortex: r(21)=0.352, p=0.099, Figure 6C & E).

#### Migraine cycle

In Migraine, lower GABA (and, consequently, GABA/Glx) levels in the thalamus were associated with being further along in the migraine cycle and subsequently, closer to the next migraine (GABA: r(15)=−0.507, p=0.038, Figure 7; GABA/Glx: r(15)=−0.559, p=0.020).

**Figure 7:**
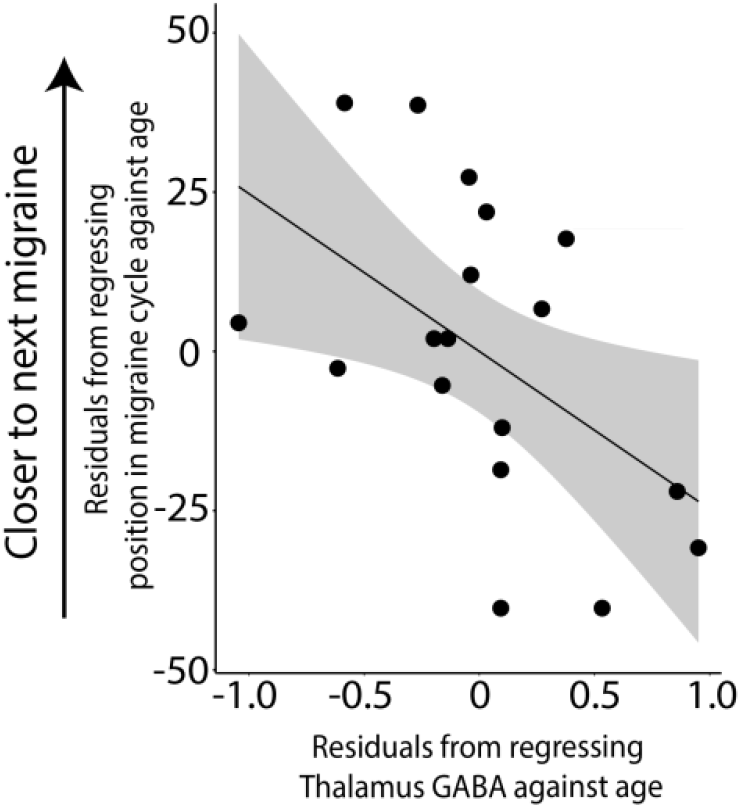
Negative correlation between levels of GABA in the thalamus and the position of the child in their migraine cycle, controlling for age (r(16)=−0.499, p=0.035).

## DISCUSSION

To our knowledge, this is the first study to measure GABA and glutamate in pediatric migraine and one of only a few studies on migraine to consider multiple voxel locations. We found (1) the migraine group had a higher GABA:Glx ratio in the thalamus as compared to the control group, which was primarily driven by an increase in GABA; (2) migraine with aura had significantly lower Glu levels in the visual cortex compared to Controls; (3) Metabolite levels in the migraine group correlate with migraine characteristics.

It has been suggested that increased glutamate is a driving force behind migraine, which is supported by the literature showing increased Glu^18,19^ or Glx^20,40^ in migraine in adults, including the analysis of subpopulations of migraine with or without aura. By contrast, we found a *decrease* in glutamate levels in the visual cortex of Migraine with aura, and a trend towards decreased glutamate in Migraine without aura compared to Controls. Aura in children is difficult to assess but is generally thought to have a similar presentation in adults and children.^4,41^ This opposite finding emphases the need to study migraine biology early. These lower glutamate levels may be a contributing factor as to why medications that are effective in adult migraineurs are not as effective in children.^42^ An early age of migraine onset is associated with an unfavourable outcome in terms of remission from migraine later in life.^45^ There is evidence that early intervention can decrease migraine frequency, with those receiving earlier interventions more likely to achieve remission^2^ Assessing early migraine biology has important implications for developing targeted, early interventions, a crucial step into reducing migraine impact throughout the lifespan.

A change from lower glutamate levels in pediatric migraine to higher glutamate levels in adult migraine may be a result of development. It is relevant to note that puberty is classed as a transitional time for migraine. Before puberty, there is a slightly higher prevalence of migraine in males, whereas after puberty prevalence increases dramatically in females, with a ratio of roughly 2 females: 1 male affected by migraine.43 The current study used a narrow age range and the majority of participants were classed as in the pre or early pubertal stage using the pubertal status questionnaire. Therefore, while it is likely that puberty has a modulatory effect on brain development and migraine, these effects are minimized here. The interaction between of puberty and changes in glutamate and GABA levels and the impact on migraine progression is important and should be answered using a longitudinal study.

In addition to higher glutamate, a recent systematic review demonstrated that adults with migraine showed higher levels of GABA in various cortical and sub-cortical regions.^46^ In the thalamus, we found GABA/Glx to be higher in Migraine, primarily driven by higher GABA, although group comparisons did not reach statistical significance. Within the migraine group, we found an association between higher GABA levels and higher migraine burden (measured by the PedMIDAS) in all three areas (although only the sensorimotor cortex reached statistical significance), indicating that children with higher GABA levels are more affected by their migraines. We also show higher glutamate in the thalamus and higher GABA/Glx ratios in the sensorimotor cortex are associated with duration since diagnosis, i.e., having migraines longer, even when controlling for current age. This suggests that these imbalances may develop over time. Indeed, there is evidence that alterations in GABA receptors influence age of onset of migraine,^47^ and adults who have suffered from migraine longer have an increased inability to habituate to stimuli.^48^ Taken together, we speculate migraine is a progressive disorder leading to more irregularities in cortical excitability with development. This highlights the potential impact of early targeted interventions on migraine progression.

To control for variations in the length of each individual’s migraine cycle, we created a novel metric to show, proportionally, how far a person was through their migraine cycle. This means the position in the cycle, along with changes in the brain associated with this, can be compared across people more accurately than simply looking at the number of days since their last migraine. We found that children in the migraine group with lower GABA levels in the thalamus were further along in their migraine cycle, and subsequently, closer to the next migraine. This suggests that, as the migraine cycle progresses, there is a reduction in inhibition in the thalamus. Evidence in adults shows an increase in cortical excitability as the migraine cycle progresses. Cortese et al. (2017)^9^ showed a negative correlation between the resting motor threshold and the time elapsed since the last attack, as the days since the last attack increased, the resting motor threshold decreased, indicating an increase in excitability in the motor cortex. Coppola et al. (2016)^10^ showed that a reduction in lateral inhibition in the somatosensory cortex was associated with a higher number of days elapsed since the last attack. These changes in cortical excitability may be driven by changes in thalamic activity or excitability over time, evidenced in the alterations in GABA levels seen here.

Our sample generally scored less than 30 on the PedMIDAS scale, indicating migraine had a mild impact on their life. While this may represent an abundance of migraine sufferers, and may reflect the typical impact of migraine in this younger sample, it is unknown whether the findings here generalize to more severe migraines that require intensive clinical management. As we see relationships between GABA levels and PedMIDAS scores, it is possible that group differences in GABA may have been detected if our migraine sample had a higher migraine burden.

In conclusion we show alterations in excitatory and inhibitory neurotransmitter levels in children with migraine, and that these measures are associated with migraine characteristics. We show higher GABA levels are associated with higher migraine burden, in line with the adult literature. We also show a reduction in glutamate levels in the visual cortex, the opposite of findings in adults. This highlights the need for further mechanistic studies of migraine in children, to aid in the development of more effective treatments.

